# Insights from controlled, comparative experiments highlight the limitations of using BSMV and FoMV for virus-enabled reverse genetics in rice

**DOI:** 10.1101/2025.08.21.671469

**Authors:** Guilherme Menegol Turra, Aldo Merotto, Dana R. MacGregor

**Affiliations:** Crop Science Department, Federal University of Rio Grande do Sul, Porto Alegre, RS, Brazil; Rothamsted Research, Harpenden, Hertfordshire, United Kingdom

**Keywords:** VIGS, VOX, VERG, *Oryza sativa*, gene silencing, overexpression

## Abstract

Virus-enabled reverse genetics (VERG) is a powerful tool for transient gene expression modulation in plants, particularly where stable transformation is challenging. However, the efficacy of VERG varies across species. In this study, we tested two commonly used viral vectors, barley stripe mosaic virus (BSMV) and foxtail mosaic virus (FoMV), for their ability to induce virus-induced gene silencing (VIGS) and virus-mediated overexpression (VOX) in rice (*Oryza sativa* L.). While both vectors successfully altered gene expression in wheat (*Triticum aestivum*), they failed to do so in six rice cultivars from different subspecies, despite rigorous optimization of inoculation methods and environmental conditions. The BSMV vector carrying antisense *phytoene desaturase* (*PDS*) sequences did not induce the expected photobleaching phenotype, and FoMV-driven GFP expression was absent in rice. These findings contrast with previous reports of successful VERG in other monocots and suggest that intrinsic resistance mechanisms exist in rice which may inhibit or reduce viral vector efficacy. Our results highlight the species-specific limitations of VERG and underscore the need for alternative viral systems or novel vector designs for functional genomics research in rice. By sharing our unsuccessful attempts, we aim to prevent redundant efforts and encourage further exploration of VERG in *Oryza* species.

## 2. Introduction

Virus-enabled reverse genetics (VERG) tools are useful for rapidly testing genotype-to-phenotype hypotheses as they transiently change gene expression *in planta*. VERG either induces or reduces gene expression, known as virus-mediated overexpression (VOX) or virus-induced gene silencing (VIGS), respectively. These techniques are particularly useful in plants where stable transformation protocols are either undeveloped or unsuitable such as for allogamous plants with long generation times and under-developed genomic resources (MacGregor, 2020). Accordingly, VERG have successfully been used in i.e. *Alopecurus myosuroides* (Mellado-Sánchez et al., 2020; Patterson et al., 2025), *Setaria italica, Solanum* spp., *Thalictrum thalictroide*, and *Zingiber officinale* (reviewed in Dommes et al., 2019). VERG are also well-established in a wide range of agronomically-important species and model plants (Dommes et al., 2019). However, not all VERG protocols are successful or repeatable, even within the same laboratory; for instance, there are conflicting reports on the efficacy of VIGS in *Pennisetum glaucum* (Schoeman, 2011; Van Nugteren et al., 2007). Its effectiveness can vary depending on host compatibility and experimental conditions (Dommes et al., 2019)

Among the most commonly used viral vectors for VERG studies, Barley stripe mosaic virus (BSMV) and Foxtail mosaic virus (FoMV) stand out due to their efficacy, flexibility and broad host ranges (Dommes et al., 2019; Rössner et al., 2022). BSMV, a tripartite, positive-sense RNA virus from the genus *Hordeivirus*, has been extensively utilized for VIGS applications in monocots, particularly in cereal crops (Lee et al., 2015a). On the other hand, FoMV, a *Potexvirus*, has demonstrated robust systemic infection in multiple monocots and dicots, making it a promising alternative to BSMV. FoMV’s relatively simple genome organization and high replication efficiency alongside a capacity to hold and retain longer inserts than BSMV contribute to its effectiveness as a VERG tool (Beernink and Whitham, 2023) and a well-characterized vector for overexpression of the GREEN FLUORECEST PROTEIN (GFP) is available (Bouton et al., 2018).

Despite the evidenced usefulness of VERG for functional genomics research, there is a distinct lack of data reporting VERG based on BSMV or FoMV in *Oryza* species, even though rice is both experimentally and agronomically important. There is evidence for rice coding sequences having been used for silencing targets in wheat driven by a BSMV-based system (Holzberg et al., 2002), for the heterologous expression of rice genes (e.g. *Hd3a*) in *Panicum miliaceum* in a FoMV-based system (Yuan et al., 2020), and more recently the use of a modified FoMV vector for silencing in japonica rice (Huang et al., 2020). Additionally, a few studies (Ding et al., 2006a, 2006b; Kant et al., 2015; Kant and Dasgupta, 2017; Purkayastha et al., 2010) have reported the use of other viral vectors in rice. Brome mosaic virus (Ding et al., 2006b) and Rice tungro bacilliform virus (Purkayastha et al., 2010) were both validated for VIGS in Asian rice (*Oryza sativa* L.) but the literature building on or effectively using their vectors is scarce. Although *O. sativa* ssp. *japonica* can be reliably transformed, efficient and effective stable transformation of *O. sativa* ssp. *indica* remains challenging (Mohammed et al., 2019) and we found no data about stable or transient transformation of weedy rice, *Oryza sativa* f. *spontanea*. Therefore, the aim of this study was to test if well-established visual reporters of VERG could induce transient gain or loss of function in *O. sativa* subspecies to gain insights in the gene regulation of this complex species.

## 3. Materials and Methods

### Plant material and growth conditions

Seeds from *Nicotiana benthamiana*, wheat (*Triticum aestivum* cv. Riband), rice (*Oryza sativa* ssp. *indica* cv. IR 64, *O. sativa* ssp. *indica* cv. Kasalath, *O. sativa* ssp. *indica* cv. CO-39, *O. sativa* ssp. *japonica* cv. Kitaake*, O. sativa* ssp. *japonica* cv. Koshihikari, *O. sativa* ssp. *japonica* cv. Balilla), and green foxtail (*Setaria viridis*) were used. All seeds except for the *S. viridis*, which were acquired from Kew’s Millenium Seed Bank (Serial Number 31491), were available from Rothamsted Research Seed Stocks. *N. benthamiana* plants were grown according (Lee et al., 2015b). All plants were grown in a high-end DEFRA/HSE-registered controlled environment room set for 26.7/21.1 ºC in a 16/8 h of light/dark regime at 200 μmol m^-2^ s^-1^. Seeds from wheat, green foxtail, and rice genotypes were surface sterilized with bleach solution (50%) for 5 minutes and rinsed 3 times with distilled water. Seeds were surfaced sowed in 20×15 cm pots containing wet Rothamsted Standard Compost Mix (75% Medium grade (L&P) peat, 12% Screened sterilised loam, 3% Medium grade vermiculite, 10% Grit (5 mm screened, lime free), 3.5 kg per m3 Osmocote® Exact Standard 3-4 M, 0.5 kg per m3 PG mix, 3kg lime pH 5.5-6.0 and 200 ml per m3 Vitax Ultrawet) and covered with perlite. The pots were covered with lids to maintain high humidity. After five to seven days, seedlings (∼2 cm) were chosen to be transplanted to square 4 cm pots filled with Rothamsted Standard Compost Mix. Propagator lids covered transplanted seedlings for 3 days following transplant. Plants were kept in the same growth room and watered every two days.

### Preparation and cloning of viral vectors

The creation of BSMV:asOsPDS1 and of BSMV:asOsPDS2 followed published procedures (Lee et al., 2015b) with minor changes detailed below. The *Oryza sativa PHYTOENE DESATURASE* (*OsPDS*, LOC_Os03g08570) was aligned with *Triticum aestivum PDS* fragments previously used (Lee et al., 2012; Mellado-Sánchez et al., 2020). Two fragments of the *OsPDS* coding sequence were synthetized in the antisense (as) direction. These covered 185 nucleotides targeting 430 to 614bp as *asOsPDS1* or 200 nucleotides targeting 900 to 1100bp as *asOsPDS2* (Figure 1). Synthetized dsDNA fragments were cloned into BSMV:RNAγ vector pCa-γbLIC (Yuan et al., 2011) and transformed into *Escherichia coli* JM109. Cloning was confirmed by colony PCR using primers in the viral backbone (2235.F - GATCAACTGCCAATCGTGAGTA and 2615.R - CCAATTCAGGCATCGTTTTC). Purified plasmid DNA from verified colonies were sequenced using the same primers to confirm cloning orientation and sequence accuracy. A selected positive transformant was transferred to *Agrobacterium tumefaciens* GV3101 following published procedures (Lee et al., 2015b). Recombinants were selected based on survival of dual selection with kanamycin and gentamycin. Individual colonies were selected, multiplied, and verified by colony PCR with the same primers described above. A selected confirmed transformant colony for each BSMV:asOsPDS1 and BSMV:asOsPDS2 was stocked in glycerol (15%) and stored at −80 ºC. BSMV:asTaPDS, BSMV:MCS, BSMV:RNAα (pCaBS-α) and BSMV:RNAβ (pCaBS-β) VIGS constructs (Lee et al., 2012; Yuan et al., 2011) and FoMV:MCS, FoMV:GFP (PV101-GFP), and P19 (pBIN61-P19) VOX constructs (Bouton et al., 2018) were used without further modifications from transformed *Agrobacterium tumefaciens* strain GV3101 stocked in 15% glycerol at −80ºC (Mellado-Sánchez et al., 2020).

**Figure 1.**
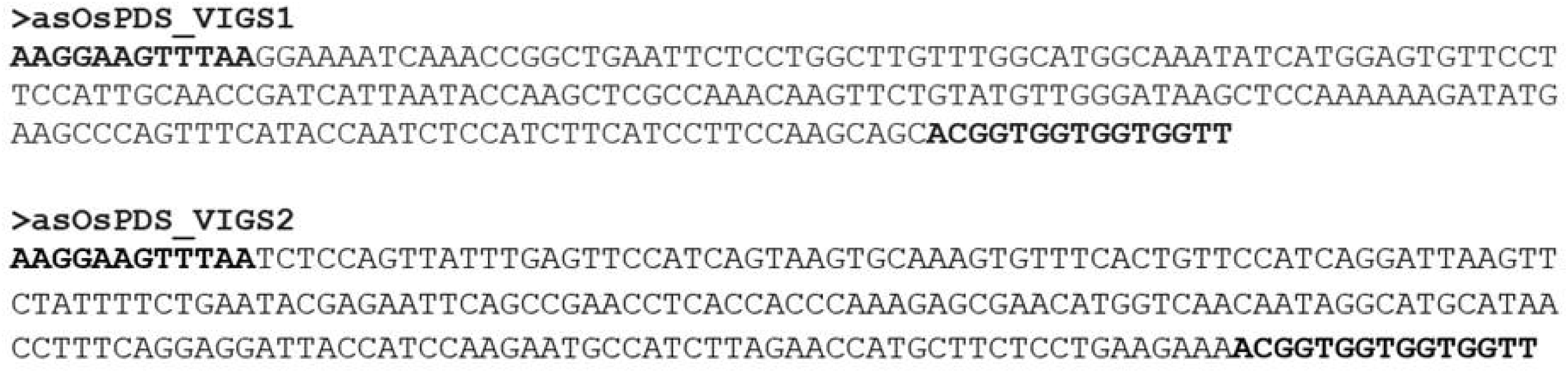
Antisense (as) Oryza sativa PDS fragments synthetised for cloning in BSMV vectors. Bold regions are ligation independent cloning adaptors.

### Preparation of the virus inoculum from Nicotiana benthamiana for rub inoculation

Detailed protocols using BSMV or FoMV are available at DOI: dx.doi.org/10.17504/protocols.io.5jyl8drr6g2w/v1. Briefly, glycerol stocks (15% glycerol with 85% bacterial culture at logarithmic growth stage) of *A. tumefaciens* containing the recombinant vectors were grown in LB broth supplemented with kanamycin (50 µg/ml) and gentamycin (25 µg/ml). A. tumefaciens cultures were pelleted and resuspended in infiltration buffer [10 mM MgCl2, 10 mM 2-(N-morpholino)ethanesulfonic acid (MES) pH 5.6, and 150 µM acetosyringone] to a final OD600 of 1.5-1.55 for BSMV constructs, 0.6 for FoMV constructs, and 0.3 for P19. BSMV:RNAγ (MCS, asTaPDS, asOsPDS1, and asOsPDS2), BSMV:RNAα, and BSMV:RNAβ were mixed in equal proportions, hereafter BSMV inoculum. FoMV:GFP and FoMV:MCS were equally mixed with P19, hereafter FoMV inoculum. BSMV and FoMV inoculums were propagated via agroinfiltration in *N. benthamiana*. Infiltrated leaves were harvested 3-5 days for BSMV and 5-7 days for FoMV after infiltration. Three leaves from different *N. benthamiana* plants were weighed into foil packets and instantly frozen in liquid nitrogen before being stored at −80°C.

### Preparation of the BSMV inoculum from Agrobacterium tumefaciens for injection inoculation and dipping

*A. tumefaciens* glycerol stocks were grown and pelleted as described above. For dipping inoculation, pelleted cultures containing BSMV constructs were resuspended in a modified dipping buffer (10 mM MgCl2, 10 mM MES pH 5.6, 200 µM acetosyringone, 1% sucrose, and 0.01% Silwet L-77) to a final OD600 of 0.6 (Andrieu et al., 2012). For injection inoculation, *A. tumefaciens* cultures were resuspended in a published injection buffer (Kant and Dasgupta, 2017) (10 mM MgCl2, 10 mM MES pH 5.6, 500 µM acetosyringone) and adjusted to a final OD600 of 0.8.

### Inoculation of plants from BSMV or FoMV infiltrated Nicotiana benthamiana

Rub-inoculation using the abrasive carborundum [Technical, SLR, Extra Fine Powder, ∼ 36 μm (300 Grit), Fisher Chemical cat 10345170] was performed as published (Lee et al., 2015b) with minor adjustments. Frozen *N. benthamiana* leaves that had been infiltrated as described above were ground in a 2:1 (w/v) ratio in 10 mM potassium phosphate buffer pH 7. The sap was used for inoculation by rubbing a leaf 6 to 10 times. Inoculation by microneedling was performed using a DermaRoller device to wound the leaf once prior the sap application. The third leaf of 14-day old wheat plants and 21-day old green foxtail and rice plants was inoculated.

### Inoculation of rice from Agrobacterium tumefaciens

For dipping inoculation, the third leaf of 21 days old rice was wounded by microneedling and dipped in the BSMV:MCS and BSMV:asOsPDS2 *A. tumefaciens* resuspensions for 30 minutes. For injection inoculation, approximately 0.2 mL of the resuspended BSMV:MCS and BSMV:asOsPDS2 inoculums were injected into the meristematic region located at the crown of 14 days old rice plants using a 1 mL syringe and a 26G needle. After inoculation, rice plants were treated in the same way as those inoculated by rubbing. Only rice cultivars IR64 and Kasalath were used for this experiment.

### Visual assessment of VIGS and VOX

Whole plants or leaf segments were photographed using a 48 megapixels camera from iPhone 15 Pro (Apple, Cupertino, CA, USA) and Velour Vinyl black backdrop (Superior Seamless 234312). For GFP fluorescence, a yellow long-pass (510 nm) filter (Midwest Optical Systems, Palatine, IL, USA) was used in front of the camera and plants were illuminated with blue light from a Dual Fluorescent Protein flashlight (Nightsea, Lexington, MA, USA).

## 4. Results and discussion

A total of 286 rice plants from subgroups *indica, aus*, and *japonica* (Table 1) were rub-inoculated. The wheat cultivar Riband was inoculated in parallel as positive control for each experiment. All experiments were conducted under controlled laboratory conditions optimized for BSMV and FoMV vectors, as temperature plays a crucial role in the effectiveness of the method (Bouton et al., 2018; Fei et al., 2021; Lee et al., 2015b). The BSMV:asOsPDS1 and BSMV:asOsPDS2 constructs created herein induced bleaching in Riband analogous to BSMV:asTaPDS, as expected (Holzberg et al., 2002), while plants inoculated with empty vector control (BSMV:MCS) remained green (Figure 2A). Systemic bleaching was seen in Riband from 12 to 35 days after inoculation (Figure 2C). Knowing sequence specificity is important for efficient VIGS (MacGregor, 2020), we used two rice-specific BSMV:asOsPDS vectors that target the same regions of PDS previously shown to be highly efficacious (Lee et al., 2015b). However, parallel inoculation of BSMV:MCS, BSMV:asOsPDS1, BSMV:asOsPDS2 or BSMV:asTaPDS into indica rice (IR64) led to a different phenotype that was similar regardless of which vector was introduced (Figure 2B). In IR64, inoculation with any BSMV vector led to development of white spots on the top of the leaf that developed after inoculation (Figure 2D). Similar white spots were also observed in uninoculated rice plants grown in the same room (Figure 2E). The lack of a clear correlation between the observed “photobleaching phenotype” and the presence of PDS sequence, or even BSMV inoculum, suggests that this phenotype is not caused by BSMV infection or target-induced VIGS but rather a consequence of the VERG growth conditions. No differences were observed between the rice genotypes tested.

**Table 1.**
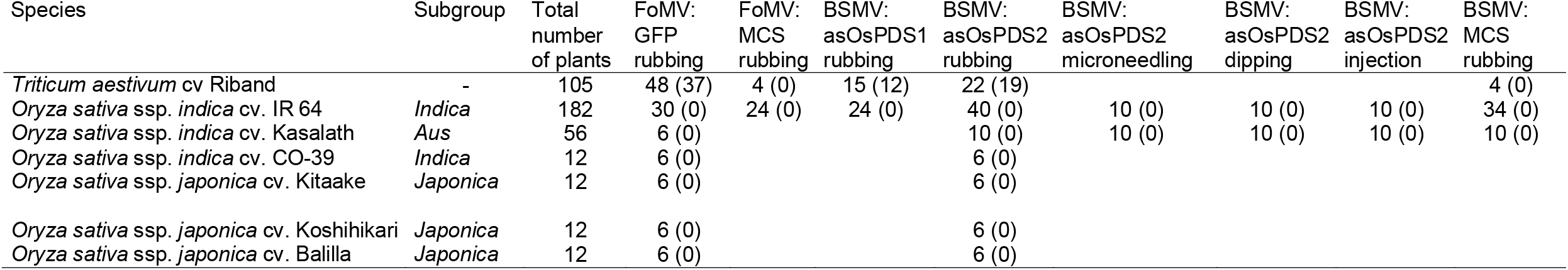
Description of plant materials and number of plants inoculated with each vector and different inoculation methods. The number in parenthesis represents plants that showed positive signs of inoculation: bleaching for BSMV and green fluorescence for FoMV:GFP

**Figure 2.**
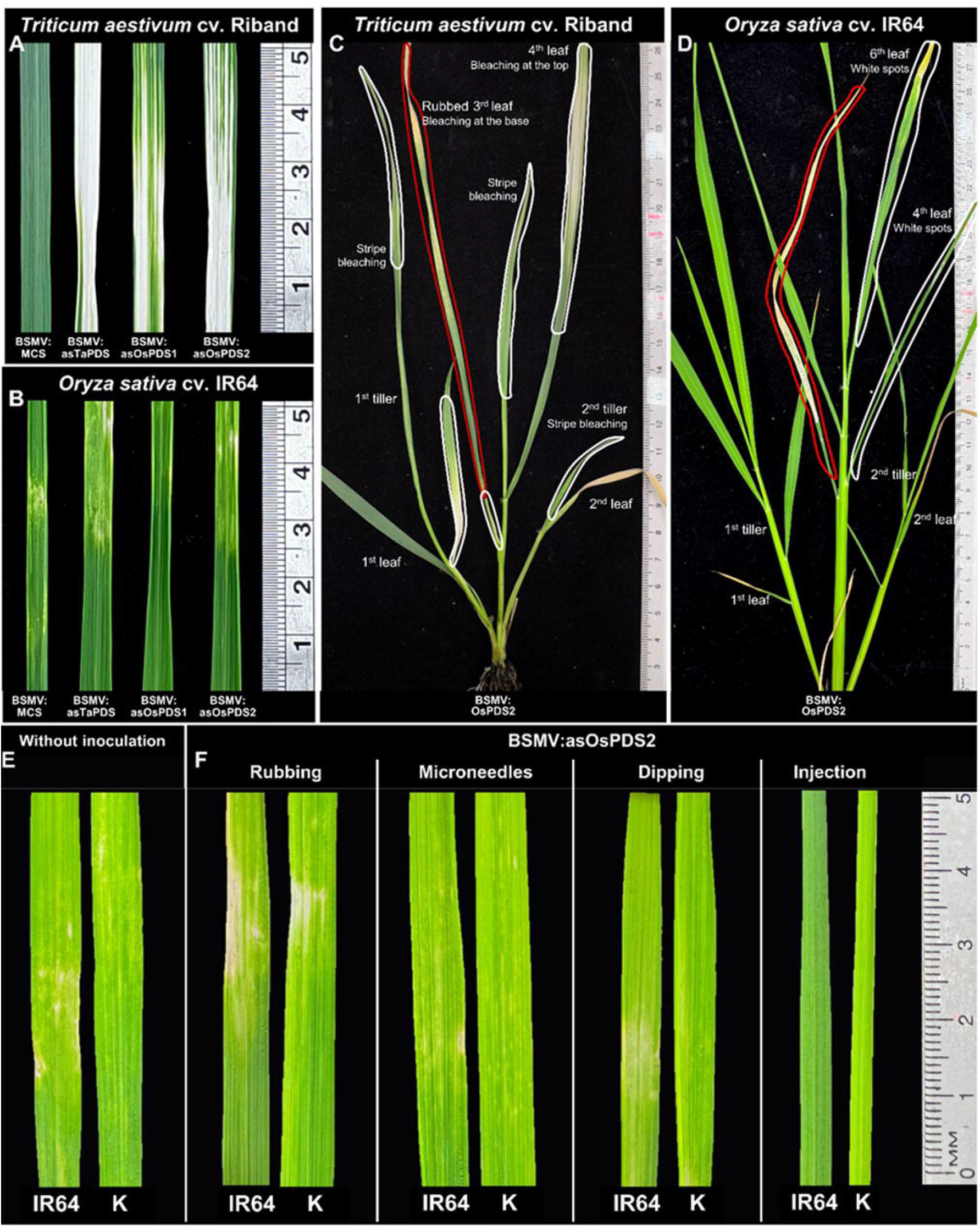
Virus-induced gene silence (VIGS) using Barley stripe mosaic virus (BSMV) vector. Phenotypes of wheat (A) and rice (B) leaves that have been inoculated with BSMV vector carrying multi cloning site (MCS) or partial sequences of *phytoene desaturase* (*PDS*) from wheat (asTaPDS) or rice (asOsPDS1 and asOsPDS2). Photos of the fourth leaf (up next the rub-inoculate leaf). Whole-plant phenotypes from wheat (C) and rice (D) that have been inoculated with BSMV:asOsPDS2. Use of different methods of inoculation were not sufficient to improve the efficacy of VIGS in *Oryza sativa* ssp. *indica* cv. IR64 from indica subgroup and *Oryza sativa* ssp. *indica* cv. Kasalath (represented by K) from aus subgroup (F). The named BSMV vector was introduced by rub-inoculation of the third leaf of 21 days old plants 6-10 times with ground leaves of *Nicotiana benthamiana* agroinfiltrated with BSMV:asOsPDS2 and the abrasive carborundum. Instead of carborundum, a device containing small metallic needles was used to wound the third leaf before the rubbing of BSMV:asOsPDS2 sap. Inoculation by dipping and injection followed published procedures with adaptations. Photos of the fourth leaf (up next the inoculated leaf) for rubbing, microneedles, and dipping and of the second leaf for injection.

In an attempt to trial many of the standard “transient expression” protocols that used rice as the target species that were available in the literature (Andrieu et al., 2012; Ding et al., 2006a, 2006b; Huang et al., 2020; Kant et al., 2015; Kant and Dasgupta, 2017; Purkayastha et al., 2010) we tried a variety of inoculation methods and resuspension buffers (see Material and Methods for details). None of these protocols (e.g. using microneedles, dipping, or injection) induced the expected photobleaching phenotype in either *indica* or *aus* rice genotypes (Figure 2F) or were more effective than the standard rub inoculation for wheat (Lee et al., 2015b). BSMV vectors driving an antisense portion of *PHYTOENE DESATURASE* are a well-accepted standard for VIGS which have been validated in several different monocot and dicot species under a range of different growth, inoculation, and propagation conditions (Lee et al., 2015b, 2012; Mellado-Sánchez et al., 2020; Patterson et al., 2025; Yuan et al., 2011). Additionally, VIGS was systematically proved to be successful in *N. benthamiana* using a wide range of heterologous inserts, as long as the viral vector is able to infect and spread (Senthil-Kumar et al., 2007). These results may indicate that there is no compatibility between BSMV and rice despite some mentions in the literature about the ability of this virus to infect the species (Benedito et al., 2004). In contrast to our data, successful VIGS was reported in rice using vectors based on Brome mosaic virus (BMV) (Ding et al., 2006b) or Rice tungro bacilliform virus (RTBV) (Purkayastha et al., 2010). There, two strains of BMV were used to generated a modified BMV vector harbouring an 86 bp fragment of the *OsPDS* used to inoculate *Oryza sativa* ssp. *indica* cv. IR64 (Ding et al., 2006b) but it is not entirely clear whether the reported phenotype is a function of *PDS* silencing or viral symptoms similar to what we observed. Similarly, RTBV was modified into a vector and tested for the ability to silence *PDS*. Its inoculation by injection in two cultivars of *indica* rice resulted in a weak streaking phenotype (Purkayastha et al., 2010). Our data also contradict the report of successful VIGS in *O. sativa* ssp. *japonica* cultivars Tatsuen No.2, Taikeng No.9, and Taoyuan No.3 using a 441 bp sequence of Barley *PDS* cloned into a modified FoMV vector (Huang et al., 2020). While their vector differed from the ones we tested, they employed a similar approach by infiltrating *Chenopodium quinoa* and rub-inoculating rice seedlings at the 2-leaf stage. We found no literature repeating or building on their successful VIGS in rice.

Inoculation with FoMV:GFP induced characteristic GFP fluorescence in Riband (Figure 3A) as expected (Bouton et al., 2018; Mei et al., 2016). Fluorescence was visible 7 to 10 days post rub-inoculation and spread systemically in Riband (Figure 3B). The same methods successfully induced GFP fluorescence in parallel in *S. viridis* (Figure 3C) and have previously been shown to be effective in *A. myosuroides* (Mellado-Sánchez et al., 2020) demonstrating that our protocols effectively and efficiently induce VOX across a variety of different monocot species. Rice plants inoculated in parallel with FoMV:MCS or FoMV:GFP showed no symptoms of viral infection or GFP fluorescence (Figure 3D) regardless rice genotype. Plants were monitored for five weeks after inoculation. FoMV driving expression of GFP (PV101-GFP) (Bouton et al., 2018) is a well-established visual reporter which has been proven to be effective in several species, particularly monocots (Bouton et al., 2018; Torti et al., 2021; Yuan et al., 2020). No reports of viral vectors that induced heterologous expression in rice have been found in the literature. In accordance to our data, FoMV has been shown to be symptomless after mechanical inoculation in rice (Paulsen, 1977). Although the efficacy of VERG systems is affected by factors such as plant growth temperature and insert orientation and length (Bouton et al., 2018; Lee et al., 2015b), ultimately, its success is dependent on the interaction between the viral vector and the host.

**Figure 3.**
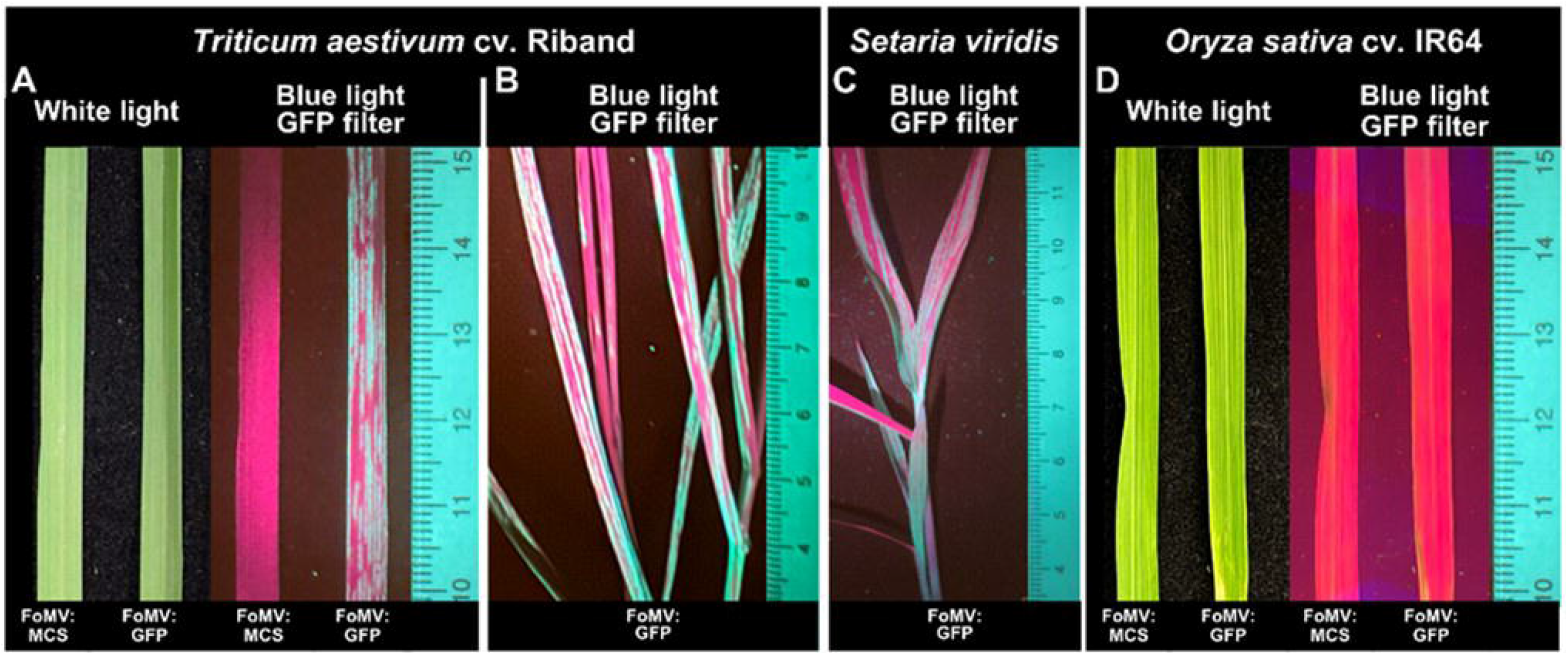
Virus-mediated protein overexpression (VOX) using Foxtail mosaic virus (FoMV) vector. Phenotypes of wheat (A) and rice (D) leaves that have been inoculated with FoMV carrying a multi cloning site (MCS) or GFP gene under white light or using blue light with a GFP filter set. Photos of the fourth leaf, up next the rub-inoculate leaf with a sap containing grinded leaves of *Nicotiana benthamiana* agroinfiltrated with FoMV:GFP and the abrasive carborundum. Phenotype of whole wheat (B) and *Setaria viridis* (C) plants that have been inoculated with FoMV:GFP showing systemic infection.

The studies reporting successful gain or loss of function in rice utilized standard growth conditions, including constant temperatures ranging from 25 to 28ºC or a 25/20ºC day/night regime, a 16/8-hour light/dark photoperiod, and relative humidity levels exceeding 80% (Ding et al., 2006a; Huang et al., 2020; Kant and Dasgupta, 2017). These conditions align with those employed in our study, suggesting that the observed ineffectiveness of the tested VERG protocols is unlikely due to environmental factors. Therefore, our data do not support the hypothesis that BSMV-driven VIGS or FoMV-driven VOX, in the form of the tested vectors, can be effectively and specifically used in the different cultivars and subspecies of *O. sativa* we tested. The possible reason for the ineffectiveness, yet to be tested, is the multiple defence pathways against viruses in rice (Qin et al., 2019). Exploring alternative viral systems or developing novel vectors specific to rice may overcome current limitations. However, as we recognise the potential impact that VERG tools could have for research in *Oryza* species, we hope that by clearly outlining our failed experimental protocols and combinations, we can help others take to alternative routes to employ VERG protocols for *O. sativa* functional genomics research.

## 5. Conclusion

This study demonstrates that barley stripe mosaic virus (BSMV) and foxtail mosaic virus (FoMV) vectors which have been optimised for virus-enabled reverse genetics (VERG) were unable to induce or reduce gene expression in rice (*O. sativa*). Although the vectors we trialled successfully altered gene expression in wheat in our experiments, or in a number of well-developed model monocots and dicots and in other non-model species, these vectors failed to induce virus-induced gene silencing (VIGS) or virus-mediated overexpression (VOX) in any of six different cultivars and subspecies of rice. Our well-controlled experiments failed despite using conditions previously optimised for viral efficacy and a variety of inoculation methods and rice genotypes. Herein, we aim to make these data public to prevent other researchers from embarking on similar attempts and to emphasize that VERG techniques, which rely on a precise balance of viral infection with plant response, are not easily transferred between species.

## 6. Acknowledgments

The authors would like to thank the Horticulture and Controlled Environment team at Rothamsted Research. The authors would like to acknowledge Margaret (Peggy) McGroary for her contributions to this research.

## 8. Data Accessibility

All data supporting the findings of this study, as well as all references to cited material, are provided within the main article. No additional datasets were generated or analysed during the current study.

## 10. Authors’ Contributions

All authors conceived the original idea and formulated the research plan; GMT designed the experiments with input from DRM; GMT performed the experiments; GMT and DRM developed the rice specific VIGS constructs; GMT wrote the manuscript with contributions from DRM and AMJ; DRM agrees to serve as the author responsible for contact and ensures communication.

